# GDNF family receptor alpha-like (GFRAL) expression is restricted to the caudal brainstem

**DOI:** 10.1101/2024.09.19.613956

**Authors:** Cecilia Hes, Luting Gui, Alexandre Bay, Fernando Alvarez, Pierce Katz, Tanushree Paul, Nadejda Bozadjieva-Kramer, Randy J. Seeley, Ciriaco A. Piccirillo, Paul Sabatini

**Author notes:** **Corresponding author:** Paul Sabatini PhD.

## Abstract

The TGF-β cytokine, growth differentiation factor 15 (GDF15) is a critical mediator of the physiologic response to a range of cellular stresses. While circulating levels of GDF15 are normally very low, these levels increase substantially under a number of acute and chronic pathogenic states including mycotoxin exposure, infection and cancer. GDF15 controls a range of physiologic outputs including reduced appetite, gastric motility, hyperalgesia, emesis, energy expenditure and immune cell function via the GDNF family receptor alpha-like (GFRAL). While the area postrema and nucleus of the solitary tract (AP/NTS) within the caudal brainstem are the only known sites of *Gfral*-expressing cells, *Gfral* may also be expressed in other cell types. We therefore utilized single molecule *in-situ* hybridizations and genetic mouse models to label *Gfral*-expressing cells from development to adult mouse. With both approaches, we found *Gfral*-labelled cells in the brainstem and extremely rare *Gfral*-labelled cells in peripheral tissues in the mouse under normal physiological conditions. Confirming these findings, single nucleus RNA-sequencing of human tissues demonstrated nearly undetectable levels of *Gfral* mRNA in sites outside the AP/NTS. Our findings confirm AP/NTS neurons are the major site of *Gfral* expression.

## 1. Introduction

GDF15 is a critical mediator of the physiologic response to stress[1]. While many cell types are capable of producing and releasing GDF15[2–4], the circulating GDF15 levels are very low under homeostatic conditions but increase under specific conditions[5]. The physiologic and pathophysiologic states in which GDF15 levels increase are highly varied and include pregnancy[2, 6], physical exercise[7], cancer[8], infection[9] and many others[10, 11]. Like the broad range of conditions which increase circulating GDF15, the functions of the hormone are equally pleiotropic. Exogenous GDF15 reduces food consumption[12], slows gastric emptying[4], and produces aversive taste responses, nausea and emesis[13]. It has also been shown to modulate the immune system[9] and increase energy expenditure[14, 15].

Given the number of tissues and cell types that express and secrete GDF15 and its wide ranging effects on different biological processes, it is somewhat unexpected that the only identified receptor for GDF15, GFRAL, is expressed exclusively in the brainstem on a small number of cells in the AP/NTS[12, 16–18]. However, it is possible that rare *Gfral*-expressing cell populations exist outside the AP/NTS. As Gfral is a lowly expressed transcript, even within the AP/NTS[19, 20], transcriptomic analysis of *Gfral* in peripheral tissues using qRT-PCR may fail to detect rare *Gfral*-expressing cells[9, 12, 16, 18, 20]. Intriguingly, a recent report details GFRAL immunoreactivity broadly across the mouse central nervous system and in multiple peripheral tissues[21], supporting a model of diffuse GDF15 action across multiple GFRAL-expressing cell types and tissues.

Thus, two models of GDF15 have emerged: one in which GDF15 action is restricted to the AP/NTS and another in which GDF15 action is mediated by multiple tissues, we sought to test these utilizing single molecule *in-situ* hybridizations (smISH), two genetic mouse models to label *Gfral*-expressing cells[19] and single-cell RNA sequencing (scRNA-seq) data[22–25].

## 2. Materials & Methods

### 2.1 Animals

Mice were bred in the Unit for Laboratory Animal Medicine at the University of Michigan and the Research Institute of McGill University Health Centre (RI-MUHC). Procedures performed were approved by the University of Michigan Committee on the Use and Care of Animals, in accordance with Association for the Assessment and Approval of Laboratory Animal Care and National Institutes of Health guidelines. Alternatively, procedures were approved by the animal care committee of McGill University. We provided mice with *ad libitum* access to food (Purina Lab Diet 5001) and water in temperature-controlled (25°C) rooms on a 12-hour light-dark cycle with daily health status checks. *Rosa26 ^LSL-eGFP-L10a^* mice[26], *Gfral^Cre^* and *Gfral^CreERT^*mice have been described previously[19]. We used both male and female mice for all studies. *Rosa26^Sun1sfGFP^* (JAX stock # 030952) mice with germline deletion of the STOP cassette were used as positive controls for green-fluorescent protein (GFP) staining.

### 2.2 Tamoxifen administration

We dissolved tamoxifen (Sigma) in corn oil and administered to mice via intraperitoneal injection once a day for 5 consecutive days at a concentration of 150mg/kg. Controls received corn oil-alone. We randomized *Gfral^CreERT^*;*Rosa26^LSL-eGFP-L10a^*mice and injected tamoxifen or a control injection. All animals were 12 weeks of age at the time of injection and tissue was collected from animals 4 weeks post-injection.

### 2.3 Tissue collection, embedding and GFP staining

We euthanized *Gfral^Cre^;Rosa26^LSL-eGFP-L10a^*mice and wild-type (WT) controls (*Gfral^WT^; Rosa26^LSL-eGFP-L10a^*) using isoflurane anaesthesia followed by CO_2_ asphyxiation. For fixed tissue collection, we perfused mice transcardially perfused with phosphate buffered saline for 3 minutes followed by 5 minutes perfusion with 10% formalin. We then collected peripheral tissues and postfixed them for 24 hours in 10% formalin before transferring them to 70% ethanol. The tissue was then dehydrated, and paraffin embedded in paraffin by the University of Michigan *in vivo* animal core. We collected, deparaffinized and rehydrated 5 µm-thick paraffin sections. We performed immunohistochemical detection of GFP with the Discovery Ultra instrument from Roche. After antigen retrieval treatment (ethylenediaminetetraacetic acid (EDTA) buffer, 32 minutes), we incubated sections 24 minutes at 37⁰C with anti-GFP (1:300, #598, MBL) followed by secondary antibody incubation with OmniMap anti-Rb HRP (760-4311, Roche) at room temperature for 20 minutes, followed by the detection Kit ChromoMap DAB kit (760-4304, Roche). Then, we counterstained the slides with the hematoxylin, and dehydrated, cleared and cover slipped them. For brain tissues, we fixed mice as described above and post-fixed brains in formalin for 24 hours prior to transferring them to 30% sucrose for a minimum of 24 hours. Subsequently, we sectioned brains as 30μm free floating sections and then blocked for 1 hour in phosphate-buffered saline with 0.1% Triton X-100 and 3% normal donkey serum (Fisher Scientific). We incubated the sections overnight at room temperature in chicken anti-GFP (GFP-1020, Aves). The following day, we washed sections and incubated them with FITC-conjugated secondary antibody (Invitrogen, Thermo Fisher, 1:300). After washing the tissues three times in phosphate-buffered saline (PBS), we mounted them onto glass slides and cover slipped them after covering in mounting media (Fluoromount-G, Southern Biotechnology). For quantification, total nuclei were quantified using Cell profiler[27] and GFP immunohistochemistry was manually counted by a blinded observer.

### 2.4 Fluorescence-activated cell sorting (FACS) for GFP+ cells

For tissue collection for FACS, we euthanized *Gfral^Cre^;Rosa26^LSL-eGFP-L10a^*mice with isoflurane followed by CO_2_ and put collected tissues in PBS. The spleen and inguinal lymph nodes were harvested separately from each mouse immediately post-mortem and kept in ∼4mL of complete RPMI 1640 (Wisent) supplemented with 10% Fetal Bovine Serum, 1% penicillin/streptomycin, 1% HEPES (Wisent), 1% sodium pyruvate (Wisent), 1% Minimum Essential Medium non-essential amino acids (Wisent), gentamycin (50mg/mL, 100µL for 500mL), and 2-mercaptoethanol. Following perfusion with complete RPMI, we mechanically dissociated the spleens and inguinal lymph nodes by gentle disruption using the plunger end of a syringe on a sterile 70μm cell strainer placed over a Petri dish. We washed cells through the strainer with 10mL of complete RPMI medium and collected them into 15mL conical tubes. We then centrifuged the resulting suspension at 400 x g for 5 minutes at 4°C to pellet the cells. We discarded the supernatant and resuspended the cell pellet in 1mL of Ammonium-Chloride-Potassium (ACK) lysis buffer (Thermofisher) and incubated for 30 seconds at room temperature to lyse erythrocytes. Lysis was terminated by quenching the suspension with 10mL of complete RPMI medium, followed by another centrifugation at 400 x g for 5 minutes at 4°C. We again discarded the supernatant and resuspended the cells in 1mL of complete RPMI medium. To ensure a single-cell suspension, we passed them through a new 70μm cell strainer. After a final centrifugation at 400 x g for 5 minutes at 4°C, we resuspended the cell pellet in 1mL of PBS. Cells were counted using a hemocytometer, and the concentration was adjusted to 1 x 10^6^ cells/mL for subsequent staining and analysis.

Cells were then incubated with Fixable Viability eFluor 780 Dye (1:1000, ThermoFisher Scientific, Cat. 65-0865-14) and Fc receptor block (1:50, 2.4G2, BD Biosciences, Cat. 553142) at 4°C for 15 min. We prepared an antibody cocktail for surface proteins in PBS and added this cocktail to the cells. We then incubated the cells at 4°C for 20 minutes before washing them with PBS and fixing them using the eBioscience Foxp3/Transcription Factor Staining Buffer Set (eBioscience). Afterwards, we washed the cells with 1X permeabilization buffer (eBioscience) and incubated them for 45 minutes at 4°C with antibody cocktails for detection of cytoplasmic and nuclear proteins. One final wash in PBS was performed before acquiring the cells on the BD LSRFortessa X-20.

We performed extracellular staining using the following antibodies (dilution, clone, company, catalog number): anti-mouse CD45.2 BUV395 (1:100, 104, BD Biosciences, Cat. 564616), anti-mouse CD3 BUV737 (1:100, 17A2, BD Biosciences, Cat. 612803), anti-mouse CD4 Alexa Fluor 700 (1:100, RM4-5, Biolegend, Cat. 100536), anti-mouse CD8α-PerCP-Cy5.5 (1:100, 53-6.7, BD Biosciences, Cat. 551162), anti-mouse CD11b V450 (1:100, M1/70, Invitrogen, Cat. 48-0112-80), anti-mouse CD19 PE (1:100, 1D3, BD Biosciences, Cat. 557399). We stained intracellular proteins using the following antibodies: anti-mouse CD3 BUV737 (1:100, 17A2, BD Biosciences, Cat. 612803) and anti-mouse CD4 Alexa Fluor 700 (1:100, RM4-5, Biolegend, Cat. 100536). We used FlowJo v10.10 software (FlowJo, LLC) to analyze data.

### 2.5 *In-situ* hybridizations (ISH)

Wild-type C57BL/6J mice (Jackson Laboratories) were euthanized and perfused with formalin as described above. We embedded kidney, pancreas, intestine, liver and ovary from each mouse in paraffin and sectioned them at 5μm thickness and fixed them on slides. We stored these slides at room temperature and then followed the ACD 323100 user manual for the RNAscope® Multiplex Fluorescent Reagent Kit v2 Assay for formalin-fixed paraffin-embedded samples. Briefly, we baked slides in a dry oven at 60°C for 1 hour after which we performed deparaffinization by incubating slides with CitriSolv (Decon Labs Inc.) twice for 5 minutes, and then with 100% ethanol twice for 2 minutes. We added H_2_O_2_ to slides and incubated them for 10 minutes while protecting them from light. In a steamer with lid, we submerged the slides in 200mL of hot diH_2_O for 10 seconds and then in 200mL RNAscope® 1X Target Retrieval Reagent for 15 minutes. After briefly transferring slides to room temperature diH2O, we submerged them in 100% ethanol for 3 minutes and let them dry. We then applied ∼1-2 drops of RNAscope® Protease Plus to each section and incubated them at 40°C for 30 minutes using a HybEZTM oven with distilled water (diH_2_O) wet paper in the tray.

In addition, we sectioned brain 30μm thick and fixed these on glass slides which were stored at -20°C for <36 hours. We then followed the ACD 323100 user manual for the RNAscope® Multiplex Fluorescent Reagent Kit v2 Assay for fixed frozen tissue samples in these sections. In this pipeline, we rinsed slides with 1X PBS and incubated them with H_2_O_2_ at room temperature for 10 minutes while protecting them from light. We removed the H_2_O_2_ and rinsed them with diH_2_O twice. In a steamer with lid, we submerged the slides in 200mL of hot diH_2_O for 10 seconds and then in 200mL RNAscope® 1X Target Retrieval Reagent for 5 minutes. After briefly transferring slides to RT diH2O, we submerged them in 100% ethanol for 3 minutes and let them dry. We then applied ∼1-2 drops of RNAscope® Protease III to each section and incubated them at 40°C for 30 minutes using a HybEZTM oven with diH_2_O wet paper in the tray.

For both brain and peripheral tissue, after rinsing the slides with diH2O, we hybridized the probes at 40°C for 2 hours. From this point on, we incubated samples at 40°C in the HybEZTM oven with humid tray and rinsed them with RNAscope® 1X Wash Buffer after each step. We followed the assay applying RNAscope® reagents AMP1, AMP2, AMP3 after which we intercalated HRP channel (15 minutes), fluorophore (30 minutes) and HRP blocker (15 minutes) for each channel in the probes mix. We used a mix of probes containing RNAscope® Mm-*Gfral* (439141-C1) and Mm-*Ppib*-C3 (312281-C3) diluted in probe diluent as specified in the manual user. The fluorophores used were Cy3 (1:1500) for *Gfral* and Cy5 (1:1000) for *Ppib,* diluted in RNAscope® TSA buffer. We applied DAPI dye (1:1000) at the end of the assay and stored samples covered from light for 48-72 hours at -4°C before imaging. Kidney, pancreas, intestine, liver and ovary were imaged on an Olympus BX61, and images of brain sections were taken on a Zeiss LSM780-NLO laser scanning confocal with IR-OPO lasers microscope at the Molecular Imaging Platform at the RI-MUHC, Montreal, CA.

### 2.6 Single-cell RNA sequencing data analysis

We used labeled human single-cell RNAseq databases[22–25] incorporated in scRNAseq v2.16.0 packages in R using SingleR v2.4.1[28]. Such databases included liver and spleen immune-resident cells[25], pancreas[23] and brain (i.e. cortex and hippocampus)[22, 24]. We used uniform manifold approximation and projection (UMAP) coordinates to visualize scaled log-normalized counts of *Gfral*. Databases were processed through Seurat v5, with libraries scaled to 10000 unique molecular identifiers (UMIs) per cell and log-normalized. We identified the most variable genes computing a bin Z-score for dispersion based on 20 bins average expression and regressed UMI counts. We then used principal component (PC) analysis for dimensionality reduction on to the top 2000 most variable genes. First, we used the first 30 PCs for UMAP projections and after integration of multiple samples contained on each database was done with Harmony[29], we used those embeddings for final UMAP projections.

## 3. Results

We followed a pipeline to interrogate several tissues for *Gfral* expression (Figure 1A). To determine if peripheral tissues expressed *Gfral* mRNA, we first performed ISH for *Gfral* on brain tissue. As a control we also probed for *Ppib*[30]. In align with previous reports[12, 17, 20], *Gfral* mRNA was readably detectable within the AP/NTS, but we failed to detect it within the hippocampus or acuate nucleus of the mediobasal hypothalamus (Figure 1B). As GFRAL-immunoreactive cells were recently described in peripheral tissues including the kidney, intestine, and liver, we also performed ISH on these tissues[21]. While *Ppib* was detected in all tissues, we failed to observe *Gfral*-expressing cells (Figure 1C).

**Figure 1.**
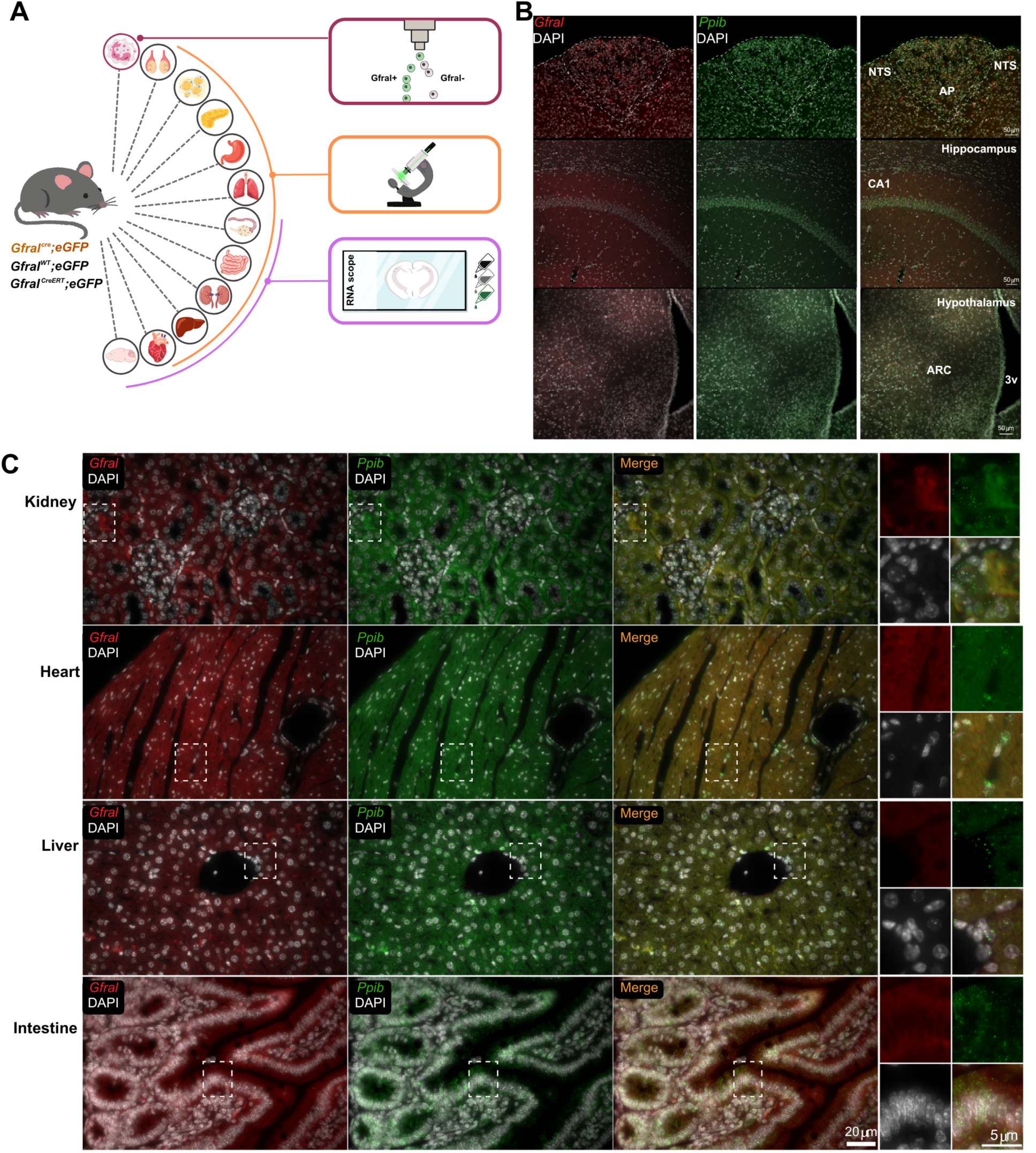
smISH for Gfral mRNA in the brain and peripheral tissues. (A) Schematic of the tissue analysis pipeline carried out in mice. We interrogated immune cells, testes, white adipose tissue, pancreas, stomach, lung, ovary, small intestine, kidney, liver, heart and brain and performed either msISH, FACS-based analysis or RNA scope *in-situ* hybridization. (B) Representative images of smISH for *Gfral* (Red, left panel) and *Ppib* mRNA (Green, center panel) and a merged image (right panel) of mouse area postrema and nucleus of the solitary tract (AP/NTS), hippocampus and medial basal hypothalamus. (C) Representative images of smISH for *Gfral* (Red) and *Ppib* mRNA (Green) with a merged image and higher magnification image of region in boxed (far right panels) of mouse kidney, heart, liver and intestine. DAPI (white) was used to stain nuclei in all images. Dashed lines in A represent the boundaries of the area postrema. **Abbreviations**: eGFP=enhanced green fluorescent protein; WT=wild-type; msISH=single molecule *in-situ* hybridization; FACS=Fluorescence activated cell sorting; AP= area postrema; NTS=nucleus of the solitary tract; ARC=arcuate nucleus; 3v=third ventricle; CA1= cornu Ammonis region 1

To complement our ISH approach to detect *Gfral*-expressing cells across a greater number of tissues, we next utilized two genetic mouse models to label *Gfral*-expressing cells with GFP. First, we crossed the *Gfral^Cre^*and Cre-dependent *eGFP-L10a* mice, generating *Gfral^Cre^*::*eGFP* mice, and examined tissues for GFP expression via immunohistochemistry (IHC) in adult *Gfral^Cre^::eGFP* mice. As the constitutively active *Gfral^Cre^* allele labels *Gfral*-expressing cells with eGFP regardless of whether *Gfral* was expressed during development or in the fully developed adult mouse, we also adopted a strategy to label *Gfral*-expressing cells exclusively in the fully developed adult mouse. This was accomplished by crossing the tamoxifen-inducible *Gfral*-cre (*Gfral^CreERT^*) with the Cre-dependent *eGFP-L10a* alleles, generating *Gfral^CreERT^::eGFP* mice. By administering tamoxifen to adult mice, we could ensure eGFP immunoreactivity (IR) was due active *Gfral* expression in developed tissues.

As expected, both *Gfral^Cre^::eGFP* and *Gfral^CreERT^::eGFP* mice displayed GFP immunoreactivity within the AP/NTS of both models, with the CreERT model displaying reduced number of GFP+ cells (Figure 2). Importantly, in the absence of Cre alleles, we failed to observe GFP expression in the *eGFP-L10* reporter (Figure 2A). We then examined multiple tissues from the *Gfral^CreERT^::eGFP* mice including heart, lung, stomach, intestine, liver, pancreas, inguinal white adipose tissue, kidney, testes and ovary (Figure 2B-C). From this analysis, we failed to detect any GFP-labelled cells in most of tissues of *Gfral^CreERT^::eGFP* mice. The exception being kidney where rare cells were detected in the renal medulla where GFP-labelled cells constitute approximately 1 of every 2000 cells (Figure 2B-C). To determine whether we could detect *Gfral*-labelled cells in the adult mouse that may have been labelled during embryonic or neonatal development, we also assayed tissues from the constitutively active *Gfral^Cre^::eGFP* mouse. Like what we observed In the *Gfral^CreERT^::eGFP* mouse, we did not detect widespread GFP-IR, rather, the only tissues we detected GFP signal was the ovary (Figure 2B-C). As we failed to detect *Gfral* mRNA by ISH in mouse ovary (Supplemental Figure 1), we assume this GFP-IR is due to developmental *Gfral* expression and indelible labelling in the adult. Importantly, a positive control for eGFP staining showed robust GFP-IR, suggesting the failure to detect GFP+ cells in the *Gfral* models was not technical in nature (Supplemental Figure 2).

**Figure 2:**
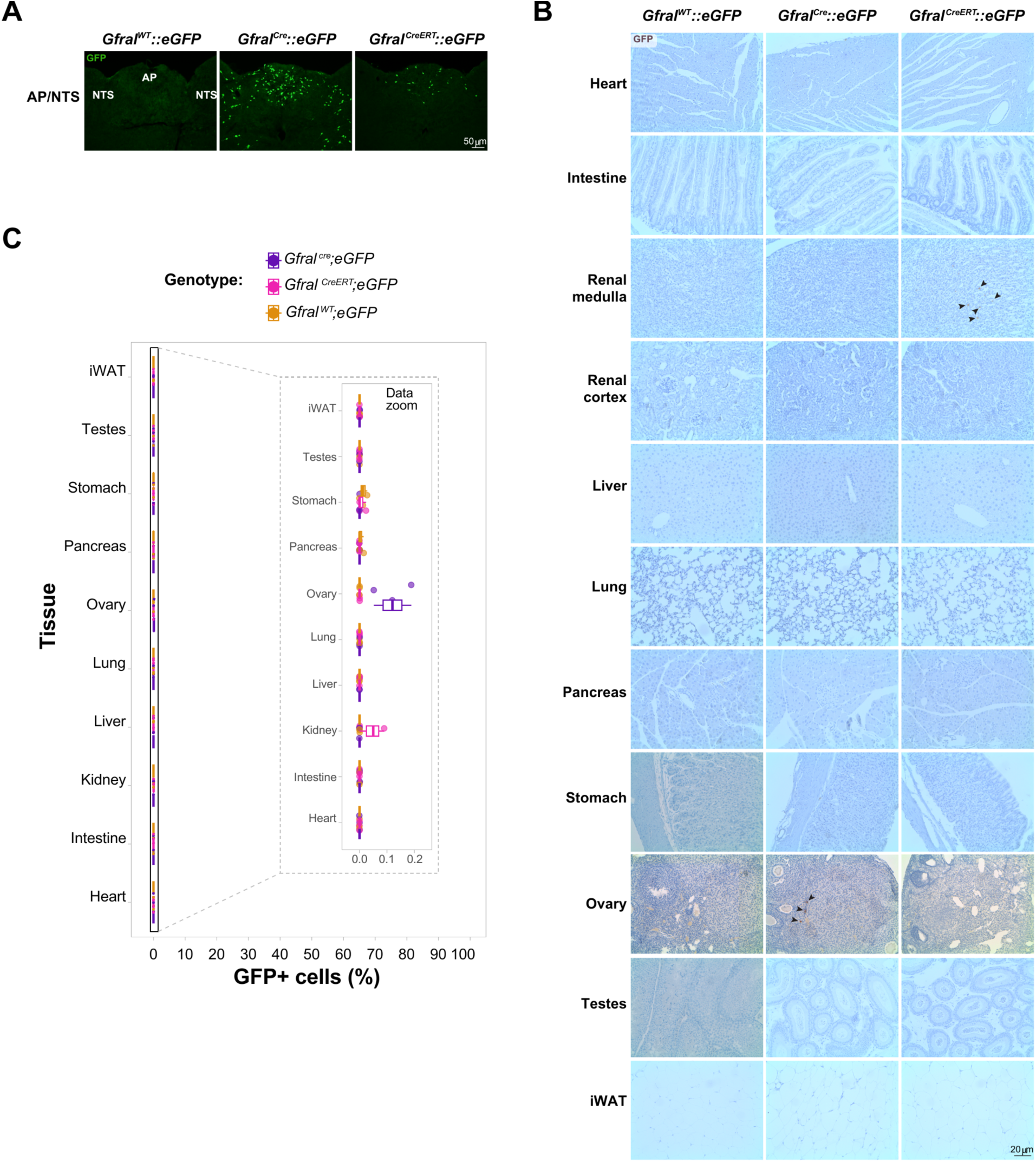
Gfral reporter expression in brainstem and peripheral tissues. (A) Representative images of GFP immunoreactivity within the AP/NTS of eGFP-L10a reporter mice lacking Cre recombinase (*Gfral^WT^::eGFP*, left panel), constitutively active *GfralCre::eGFP* (middle panel) and tamoxifen inducible *Gfral^CreERT^::eGFP* mice (right panel). (B) Representative images of GFP immunoreactivity assessed through immunohistochemistry for GFP-labelled cells in mouse tissues from *Gfral^WT^::eGFP* (left panels), *GfralCre::eGFP* (center panels), *Gfral^CreERT^::eGFP* (right panels) animals. (C) Combined boxplot and dotplot of the quantification of GFP positive cells in AP/NTS and peripheral tissues. Three samples per tissue per genotype were quantified. The percentage of cells are given regarding total nuclei on each sample. **Abbreviations:** eGFP=enhanced green fluorescent protein; GFP=enhanced green fluorescent protein; AP= area postrema; NTS=nucleus of the solitary tract; iWAT=inguinal white adipose tissue; WT=wild-type

As GDF15 has roles regulating the function of numerous immune cell types[31, 32], we evaluated GFP expression in various immune cell types from secondary lymphoid tissues (spleen and L.N.s) from *Gfral^Cre^::eGFP* mice (Figure 3). Our results show that the frequency or level (Figure 3B-C) of GFP expression in total hematopoietic (CD45+), T (CD3+), B (CD19+) and myeloid (CD11b+) cells in spleen is not significantly different between mice of both genotypes. Interestingly, within the myeloid compartment of the inguinal lymph node, we found a higher number of GFP cells. However, when normalized to total myeloid cell counts, the number of GFP+ cells was not significantly higher than GFP+ counts from control animals. Collectively, these results suggest *Gfral* is not expressed in immune cells from healthy mice.

**Figure 3:**
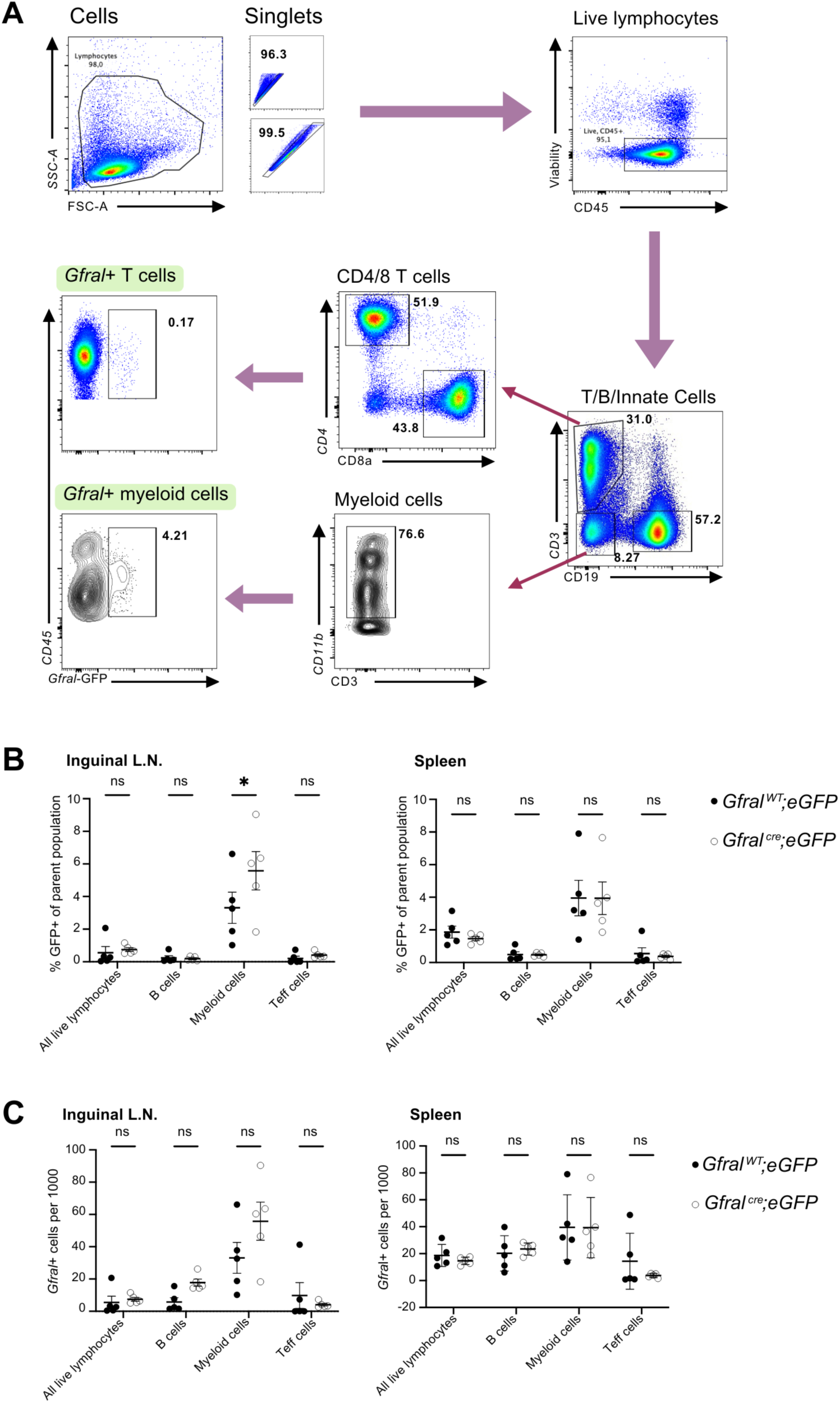
Gfral Cre reporter expression in immune cells. (A) Representative flow cytometry analysis of GFP reporter expression in CD45+ immune cells, further sub-gated by differential expression of CD4, CD8, CD11b, and CD19. (B) Mean frequency of GFP+ cells in the pre-gated parent population of cells taken from the inguinal lymph nodes (left) and spleen (right) (n=5) compared to control (n=5). (C) Mean count of GFP+ cells adjusted per 1000 cells in the pre-gated parent population of cells taken from the inguinal lymph nodes (left) and spleen (right) (n=5) compared to controls (n=5). Numbers in FACS plots indicate the frequencies of gated cells. Error bars, SEM. *p-value≤0.05 determined by Two-Way ANOVA. **Abbreviations:** eGFP=enhanced green fluorescent protein; GFP=enhanced green fluorescent protein; WT=wild-type; ns=non-significant; L.N.=lymph node; FACS=Fluorescence activated cell sorting; SEM=standard error of the mean

Finally, we made use of publicly available single cell RNA-sequencing (scRNA-seq) labeled datasets[22, 23, 25] to corroborate our results in human adult and embryonic samples[24] (Figure 4). Except for rare cell expression in cortex and hippocampus, in human pancreas and brain, we were unable to find *Gfral* expressing cells (Figure 4A). The rare brain expression was not associated with developmental stages or age (Supplemental Figure 3).

**Figure 4:**
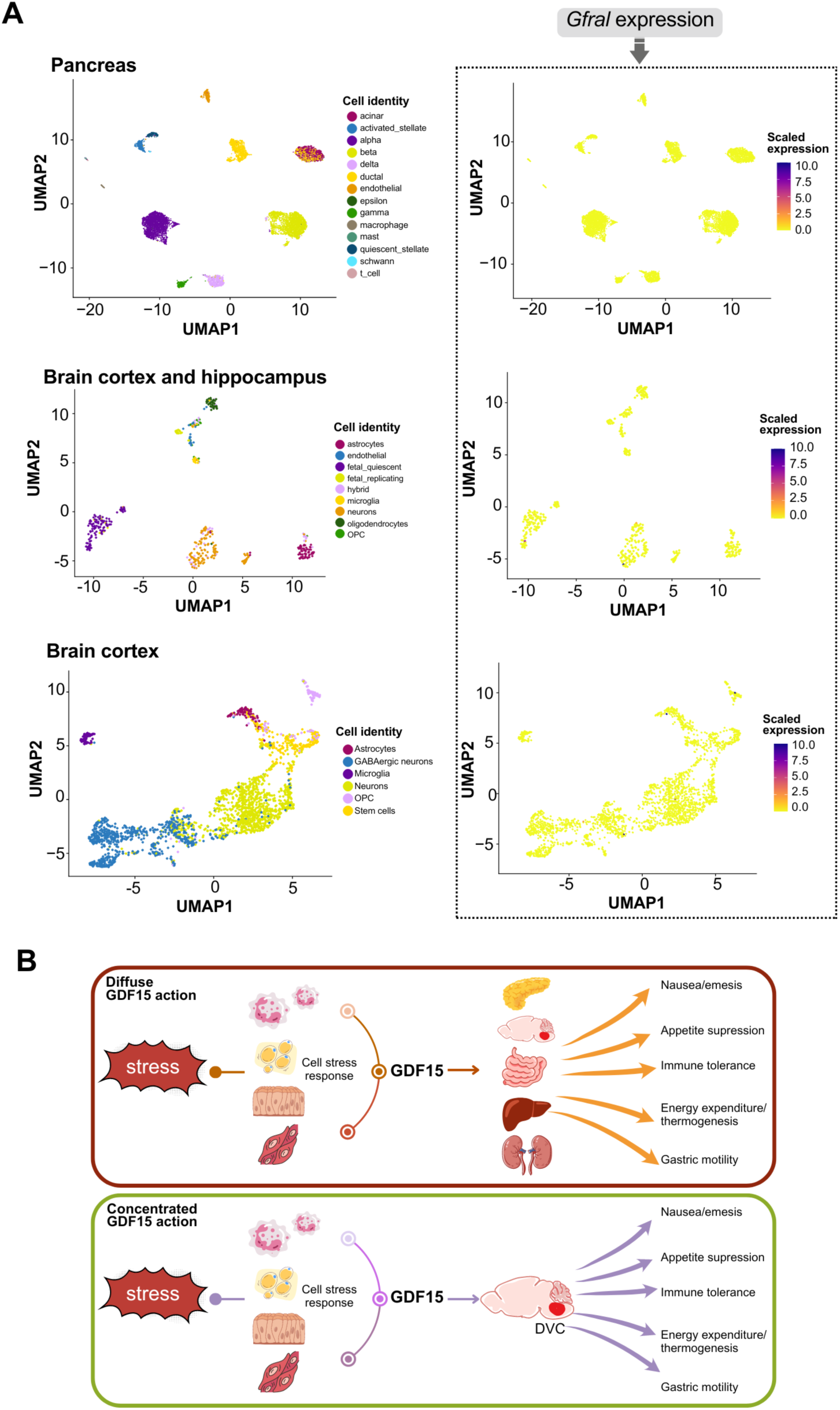
Gfral expression in human tissues using scRNA-seq data. (A) UMAP plot of the scRNA-seq publicly available data on adult and developmental tissues showing *Gfral* expression (right panels) on each cell type (left panel). (B) Models of the GDF15/GFRAL interaction proposed. The DVC includes the area postrema and the nucleus of the solitary tract in the hindbrain. **Abbreviations:** UMAP= uniform manifold approximation and projection; scRNA-seq= single-cell RNA-sequencing; OPC=oligodendrocyte precursor cell; GABA= gamma-aminobutyric acid; DVC=dorsal vagal complex

Overall, our results show a conserved and restricted expression of GFRAL in the AP/NTS supporting a concentrated, rather than a diffuse, interaction with GDF15 to mediate this cytokine effects (Figure 4B).

## 4. Discussion

In mammalian physiology individual peptide hormones are generally expressed by a small number of specialized cells within a single tissue and the respective receptor(s) are expressed broadly across many cell types and tissues. Such is the case with insulin, glucagon, growth hormone and leptin[33–36]. However, the GDF15-GFRAL system seems to function uniquely as many different tissues and cell types can produce GDF15 but the expression of GFRAL, the only well-described receptor for GDF15, seems to be highly restricted to a single site across the entire body. GDF15 is the only known ligand for GFRAL and it regulates a range of physiologies including appetite, gastric motility, nausea and emesis, immune cell function and energy expenditure [4, 9, 12–15]. One possible mechanism to explain how GDF15 regulates such variate range of physiologic roles is that all these actions are solely mediated via GFRAL signaling on AP and NTS neurons, but a separate model could be that GDF15 acts on GFRAL expressing cells on peripheral tissues. To date, most reports support the former model with multiple analysis of *Gfral* mRNA using highly sensitive approaches failing to detect *Gfral* mRNA in brain regions outside the AP/NTS[12, 16–18, 20].

Within the periphery, experiments using transcriptomic approaches to measure mRNA such as qRT-PCR, have not found appreciable levels of *Gfral* in many peripheral tissues[9, 12, 16, 18, 20]. However, this does not rule out the existence of rare *Gfral-* expressing cell populations that mediate some of GDF15 effects in regions other than the AP/NTS. Indeed, a recent study observed GFRAL immunoreactivity in multiple peripheral tissues and even within central nervous system sites beyond the AP/NTS[21], raising the possibility for the existence of uncharacterized GFRAL-expressing cells which would have direct implications on the development of GDF15/GFRAL-based therapies and impair the therapeutic advantage of GFRAL restricted expression limiting off-target complications. The antibody used in these studies has previously been shown to be specific for GFRAL within the AP/NTS[20] and there is a lack of validation in other central nervous system sites and peripheral tissues where it may bind other proteins and result in a false positive. Due to this, we sought to determine whether *Gfral* is expressed in peripheral tissues using three approaches.

First, smISH which is specific for targeted sequences, failed to detect *Gfral* mRNA in any mouse peripheral tissues we tested. Additionally, we used two genetic models to label *Gfral-*expressing cells with GFP either throughout development (*Gfral^Cre^::eGFP*) or specifically in the adult (*Gfral^CreERT^::eGFP*). Importantly, these genetic models have demonstrated a high concordance between Cre recombinase and *Gfral* using smFISH and translating ribosomal affinity purification (TRAP)[19]. Screening peripheral tissues for GFP-labelled cells, revealed rare GFP-labelled cells in kidney, pancreas and ovary. Of these tissues, GFP was also detected in the pancreas in the absence of Cre recombinase, suggesting a degree of “leak” in the Cre-dependent reporter depending on the tissue. Separately, we noted higher numbers of GFP+ myeloid cells of the lymph node, however this increase was not significantly higher than the number of GFP+ cells in control animals, likely due to autofluorescence. To contrast the mouse experiments with human tissues, we examined immune-resident cells of the liver and the spleen from human tissue[25] but as the *GFRAL* gene was not mapped due to a complete lack of reads mapping to this gene. Therefore, within the datasets we could not acquire a plot or quantification of expression and further suggesting that *GFRAL* is not expressed by immune cells.

Additionally, we examined single cell sequencing data from five human labeled datasets of non-brainstem cells including pancreas, brain cortex, hippocampus and immune-resident cells of the liver and the spleen[22–24]. Similar to what we observed in the mouse, *GFRAL*-expressing cells in human tissues were undetectable in most analyzed tissues. In cortex and hippocampus *Gfral*+ cells could be detected regardless of developmental stage but were extremely rare. This supports our proposed model of concentrated GDF15 action mediated via GFRAL expression in the AP/NTS.

Limitations of our study include that we assume any *Gfral^Cre^*-expressing cell will also express GFP. However, this is not always the case as Cre-reporters do not fully represent their Cre-driver[37] However, the addition the lack of *Gfral* mRNA detected by *in-situ* hybridizations in these same tissues gives us confidence in our genetic approach to identify *Gfral*-expressing cells. Additionally, while we attempted to examine a broad range of tissue types, there are tissues and cell types not examined herein and these may warrant further investigation. Finally, regardless of other receptors besides GFRAL for GDF15 action have been proposed by some reports[38, 39], our study does not address GDF15 interaction with peripheral tissues via other receptors. For instance, we proved that *Gfral* is not expressed in immune cells in healthy mice and human immune cells spleen and liver-resident, and this may be due the expression of another receptor in these cells to be modulated by GDF15 as has been reported in T regulatory cells (i.e. CD48)[38].

## 5. Conclusion

GDF15-GFRAL signaling is a critical mediator of the physiologic response to stress and activates numerous pathways that reduce appetite, alter gut motility, promote nausea, emesis and taste aversions while also increasing energy expenditure and regulating the immune system. As these pleiotropic effects could be mediated widespread GFRAL expression, we sought to quantify *Gfral*-labelled cells in multiple mouse and human tissues using highly sensitive and specific approaches. While *Gfral*-labelled cells could be detected in some tissues including mouse ovary and kidney, these cells were extraordinarily rare and their biologic relevance, uncertain. Together, our data supports the model wherein GDF15 action is largely, if not exclusively, mediated by GFRAL expressed within the AP/NTS.

## 6. Acknowledgements

We acknowledge the technical assistance of the University of Michigan and the Histopathology core of the Research institute of the McGill University health centre. We also thank F. Soltani for technical assistance for ISH.

## 7. Author contributions

Conceptualization: PS, RJS and CAP

Data curation: CH, AB, FA, TP and PS

Formal analysis: CH, AB, FA, PK and PS

Funding acquisition: PS

Investigation: CH, LG, AB, FA, PK, TP, NBK and PS

Methodology: CH, RJS, CAP and PS

Sample collection: LG and PS

Software: CH, AB and FA

Visualization: CH, AB, FA and PS

Writing—original draft: CH, AB and PS —review and editing: CH, LG, AB, FA, PK, TP, NBK, RJS, CAP and PS

