## Supplementary figures and images for "GDNF family receptor alpha-like (GFRAL) expression is restricted to the caudal brainstem"

### Supplemental figures

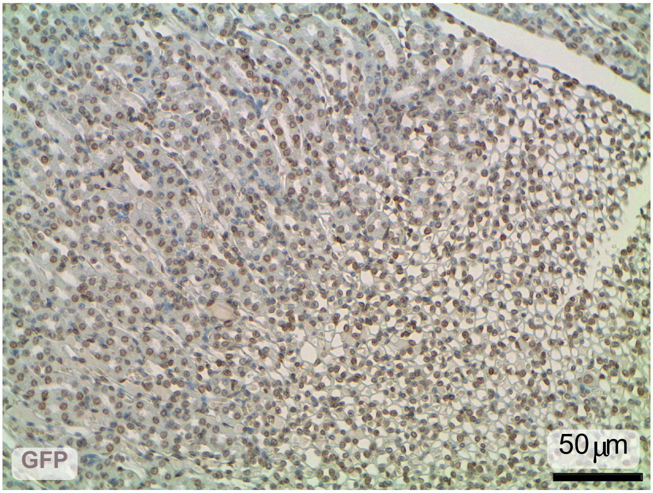


**Supplemental Figure 1**

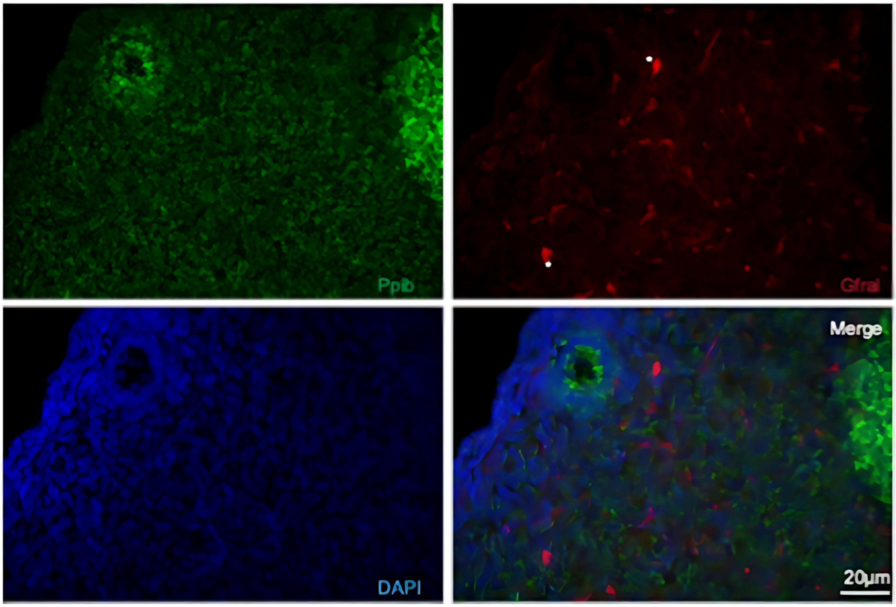


Supplemental Figure 2

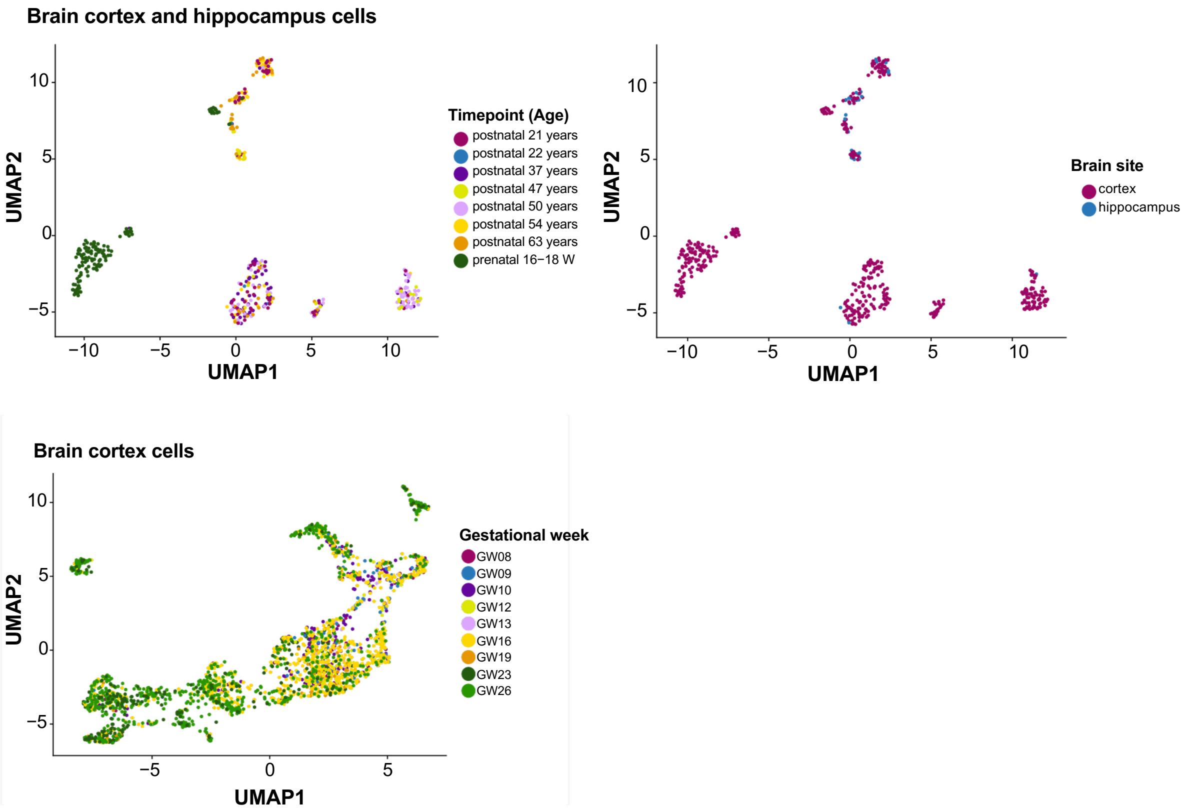


**Supplemental Figure 3**
